# Efficient derivation and transcriptional characterization of mouse extra-embryonic endoderm stem cell lines generated by somatic cell nuclear transfer

**DOI:** 10.64898/2026.02.22.707260

**Authors:** Shuaipeng Li, Shu Wei, Guomeng Li, Mei Hu, Jiangwei Lin, Wandong Bao

**Affiliations:** Department of Laboratory Animal Science, Kunming Medical University, Kunming, Yunnan, 650500, China; State Key Laboratory of Genetic Evolution & Animal Models, Kunming Institute of Zoology, Chinese Academy of Sciences, Kunming, Yunnan, 650223, China

**Author notes:** Corresponding author. Correspondence: Jiangwei Lin,; Wandong Bao,. Contributed equally (Shuaipeng Li, Shu Wei).

**Keywords:** Somatic cell nuclear transfer, Primitive endoderm, Extra-embryonic endoderm stem cell line, NanoString multiplex gene expression profiling, Aberrant epigenetic reprogramming, Imprinting errors

## Abstract

Somatic cell nuclear transfer (SCNT) holds great promise for regenerative medicine and agriculture, but its application is severely hampered by low efficiency, primarily attributable to aberrant epigenetic reprogramming. Although embryonic stem cells (ESCs) and trophoblast stem cells (TSCs) have been successfully derived from cloned embryos, an *in vitro* counterpart of the primitive endoderm (PrE) lineage has remained unavailable. To address this gap, this study reports the first successful establishment of extra-embryonic endoderm stem cell lines (XENs) from mouse SCNT-derived blastocysts (NT-XENs). Under conventional culture conditions, NT-XENs were generated from hybrid B6D2F1 blastocysts at a high efficiency of 55%, comparable to that of fertilization-derived XEN lines (FD-XENs, 50%), whereas derivation from inbred C57BL/6J SCNT-derived blastocysts was markedly lower (12.5%). Immunofluorescence and NanoString multiplex gene expression profiling confirmed that NT-XENs robustly expressed specific marker genes for PrE/XENs (e.g., *Gata4, Gata6, Sox17*), while exhibiting negligible or absent expression of pluripotency and trophoblast markers. Based on NanoString assay data, NT-XENs and FD-XENs shared highly similar global gene expression patterns, yet also exhibited some nonnegligible differences, exemplified by the differentially expressed genes (DEGs) *Pecam1, Gtl2, Thbd* and *Xlr3b*, which may suggest that the NT-XENs resided in a more differentiated state (potentially biased toward parietal endoderm (PE)) and retained SCNT-specific epigenetic imprinting errors, including aberrant X-chromosome inactivation and dysregulation of imprinted domains. In summary, this study successfully establishes NT-XEN cell lines, providing a valuable *in vitro* model for investigating the reprogramming scenarios of PrE lineage in SCNT and offering novel insights into the mechanisms underlying developmental failure of cloned embryos.

## Introduction

SCNT is a revolutionary technology that enables the production of genetically identical individuals from differentiated somatic cells, holding immense promise for a range of applications in agriculture and regenerative medicine [1]. Despite its significant potential, the widespread application of SCNT is severely impeded by the remarkably low efficiency, i.e. the vast majority of cloned embryos fail to develop to term [2]. This high rate of developmental failure is widely attributed to incomplete or erroneous epigenetic reprogramming of the donor somatic nucleus [3], of which the DNA methylation and histone modification profiles must be comprehensively remodeled into a totipotent embryonic state for achieving cloned animals [4]. A substantial body of evidences indicate that these reprogramming errors disproportionately compromise the development of extra-embryonic tissues [5]. Specifically, abnormalities such as placental insufficiency and defective yolk sac formation are frequently observed in cloned conceptuses and are considered major contributing factors to their demise [6].

The formation of these critical supportive tissues is rooted in a series of precisely orchestrated cell fate decisions during early mammalian embryonic development. In the blastocyst, the first lineage segregation event establishes the trophectoderm (TE) and the inner cell mass (ICM) [7, 8]. The ICM subsequently undergoes a second lineage segregation, differentiating into the pluripotent epiblast (EPI), the source of the fetus proper, and the PrE [9]. The PrE lineage further differentiates into two distinct layers, the PE and the visceral endoderm (VE) [10, 11], which together construct the yolk sac indispensable for providing crucial nutritional and signaling support to the nascent embryo before the placenta becomes fully functional [12, 13]. The coordinated development and interaction of EPI, PrE and TE are critical for embryonic viability, and their corresponding self-renewing stem cell lines have been derived *in vitro*, respectively: ESCs from the EPI [14], XENs from the PrE [15], and TSCs from the TE [16]. The availability of these stem cell lines provides a powerful and tractable toolkit for dissecting the molecular underpinnings of lineage specification and function.

Among these cellular models, XENs offer a unique *in vitro* system to study PrE-associated biological mechanisms. They are robustly characterized by the expression of a suite of key transcription factors, such as GATA4, GATA6, SOX7, and SOX17, as well as the cell surface receptor PDGFRA [15, 17]. While both nuclear transfer-derived ESCs (NT-ESCs) [18, 19] and nuclear transfer-derived TSCs (NT-TSCs) [20] have been successfully established from SCNT-derived embryos, offering invaluable insights into the reprogramming fidelity of the EPI and TE lineages, a stable *in vitro* model for the PrE lineage originating from cloned embryos has been conspicuously absent. To address this gap, our study reports the first successful derivation, with a remarkable efficiency, and characterization of the XEN cell lines from SCNT-derived blastocysts. Utilizing the targeted analysis of key developmental genes, we profiled the transcriptional state of the NT-XENs and examined their distinctive features from the FD-XENs.

## Materials and methods

### Mouse strains

The mice of wild-type C57BL/6J and DBA/2 genetic background used in the present study were purchased from Charles River Biotechnology Co. LTD (Beijing, China). Mouse care and all the experimental procedures were conducted in compliance with the guidelines of the Institutional Animal Care and Use Committee (IACUC) of the Kunming Institute of Zoology, Chinese Academy of Sciences. The approval number for all the contents of this research is IACUC-RE-2024-01-006.

### Cell culture medium

DMEM was supplemented with 15% FBS, 1% penicillin/streptomycin, 0.1 mM nonessential amino acids, 1% β-mercaptoethanol, 2 mM GlutaMAX Supplement, 1 mM sodium pyruvate, and 1,000 IU/mL LIF. This ES medium with LIF was used for deriving XENs from blastocysts.

### Nuclear transfer

Metaphase II-arrested oocytes were collected from superovulated B6D2F1 females (8-10 weeks, generated by mating female C57BL/6J with male DBA/2) and cumulus cells were removed using hyaluronidase. The oocytes were enucleated in a droplet of HEPES-CZB medium containing 5 µg/mL cytochalasin B (CB) using a blunt Piezo-driven pipette. Then, the spindle-free oocytes were washed extensively and maintained in CZB medium up to 2 hr before nucleus injection. The cumulus cells from B6D2F1 females were gently aspirated in and out of the injection pipette to remove the cytoplasmic material and then injected into enucleated oocytes. The reconstructed oocytes were cultured in CZB medium for 1 hr and then activated for 5-6 hr in activation medium containing 10 mM Sr^2+^, 5 ng/mL trichostatin A (TSA) and 5 µg/mL CB. All of the reconstructed embryos were subsequently cultured in KSOM medium supplemented with 5 ng/mL TSA for another 3-4 hr, and then maintained in KSOM medium with amino acids at 37 □ under 5% CO_2_ in air. For the nuclear transfer in an inbred strain background, metaphase II-arrested oocytes and cumulus cells from wildtype C57BL/6J females were used, with all other experimental procedures above unchanged.

### Derivation of XEN cell lines from blastocysts

The SCNT-derived embryos were cultured in KSOM medium with amino acids to the blastocyst stage, so were the B6D2F1 fertilization-derived embryos (generated by mating female C57BL/6J with male DBA/2) after collection at the 2- to 8-cell stage. Upon reaching the blastocyst stage, the zona pellucida was dissolved by exposure to acid Tyrode’s solution. Subsequently, individual blastocysts were plated onto 0.1% gelatin-coated 4-well plates containing a feeder layer of MEFs. Culture was maintained using ES medium containing LIF to promote XEN cell derivation, following previously established protocols [21].

### Immunofluorescence and imaging

Immunofluorescence staining was performed as previously described [22]. Briefly, cell lines were cultured in 4- or 24-well dishes. Cells were fixed in 4% paraformaldehyde at 4 □ overnight or room temperature for 30 min, permeabilized with 0.1% Triton X-100 in 1× PBS (1× PBST) for 30 min, and blocked with 5% normal donkey serum diluted in 1× PBST (blocking solution) for 1 hr. Primary antibodies were diluted at 1:50–1:200 in blocking solution and samples were then incubated at 4 □ with rotation overnight. After three 10-min washes in 1× PBST, samples were incubated in a 1:500 dilution of secondary antibody in blocking solution for 1–1.5 hr at room temperature, then washed and covered with 1× PBST containing DAPI. Images were taken with an AMG EVOS fluorescence microscope (Life Technologies). Primary antibodies from Santa Cruz Biotechnology were against Gata4 (cat. # SC-1237), Dab2 (cat. # SC-13982), Oct4 (cat. # SC-5279), Nanog (cat. # SC-376915), and Cdx2 (cat. # SC-166830). Primary antibodies from R&D Systems were against Gata6 (cat. # AF1700) and Sox17 (cat. # AF1924). Appropriate secondary antibodies were purchased from Jackson ImmunoResearch Laboratories and Invitrogen.

### NanoString multiplex gene expression analysis

NanoString assays were performed as previously described [22] with necessary modifications for this research. Briefly, dissociated cells were collected by trypsinization and centrifugation. Cell pellets were dispensed in RNAlater Stabilization Solution (Qiagen) and stored at -80 □ for later use. Cell pellets were lysed in RLT Lysis Plus Buffer using a TissueLyser LT (Qiagen) at 40 Hz for 2 min, then total RNA samples were extracted using RNeasy Plus Micro kit (Qiagen) according to manufacturer’s protocol. The custom NanoString CodeSet “Extra” was used. For each total RNA sample, an aliquot of 100 ng was hybridized at 65 □ for 18 hr and processed with the nCounter Analysis System GEN1 (NanoString Technologies). The reporter counts were processed using nSolver Analysis Software v2.5 (NanoString). Two normalizations were performed to the counts, the first normalization to the generic positive controls, followed by normalization to the reference genes, *Pgk1* and *Hsp90ab1*. The normalized counts were then log2-transformed, and used for the correlation clustering analysis and heatmap plotting of stem cell line-specific marker gene expressions across all samples. The DEG analysis between NT-XENs and FD-XENs was performed using the limma R package.

### Figure preparation

The experimental workflow diagram shown in Figure 1A was created using the graphical assets from BioGDP.com [23]. Multichannel-merged immunofluorescence images were created using ImageJ. All NanoString assay data-associated figures in Figure 3 were plotted using the Metware Cloud (https://cloud.metware.cn).

**Figure 1.**
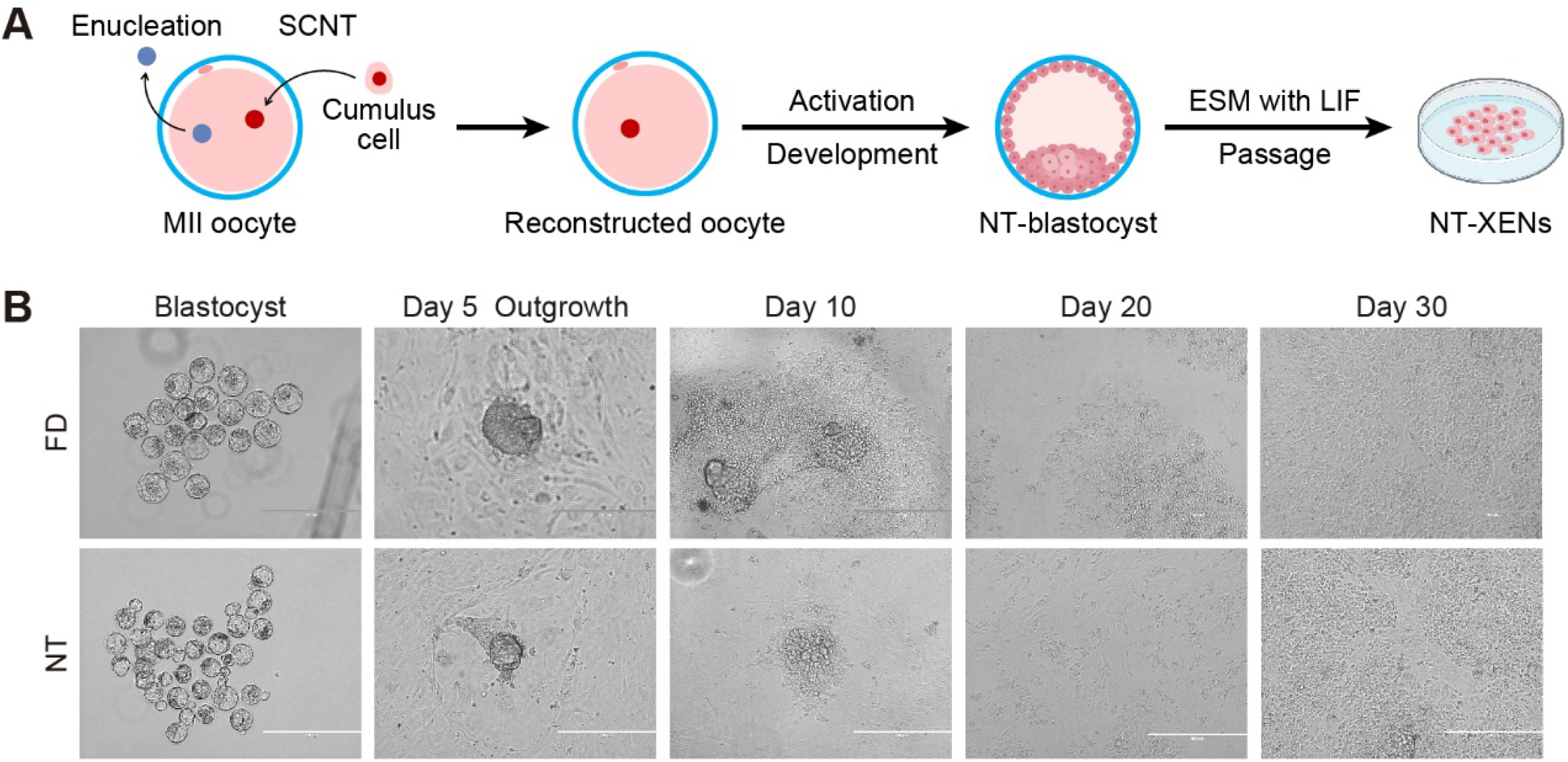
Derivation of XENs from mouse SCNT-derived blastocysts. (A) Experimental strategy for establishing mouse nuclear transfer-derived XEN cell lines. MII oocytes were enucleated and received nuclear transfer from cumulus cells to form reconstructed oocytes, which were subsequently activated and developed to the blastocyst stage *in vitro*. These SCNT-derived blastocysts were then cultured in ES medium with LIF to derive XEN cells, and the NT-XEN cell lines were ultimately established via continuous subculture. MII, metaphase II; SCNT, somatic cell nuclear transfer; ESM, ES medium; LIF, leukemia inhibitory factor. (B) The process of deriving XEN cells, respectively from mouse fertilization-derived and SCNT-derived blastocysts, and establishing their cell lines *in vitro* under bright-field microscopy. The numbers of days cultured in ES medium with LIF are indicated. Scale bar: 400 μm.

## Results

### Efficient derivation of XEN cell lines from SCNT-derived blastocysts

To derive NT-XEN cell lines, we reconstructed embryos by enucleating the metaphase II (MII) oocytes from hybrid B6D2F1 females (generated by mating female C57BL/6J with male DBA/2) and injecting the nuclei from their own cumulus cells. The reconstructed embryos were cultured to the blastocyst stage *in vitro* and then used to establish XEN lines in ES medium supplemented with leukemia inhibitory factor (LIF) (Figure 1A). As a control, fertilization-derived blastocysts of B6D2F1 background were also processed under identical culture conditions for XEN line derivation, after their *in vitro* development from the early cleavage stage.

Specifically, we cultured the reconstructed embryos in KSOM medium until the blastocyst stage and then removed the zona pellucida. Radom 20 of these SCNT-derived blastocysts were separately seeded into the wells coated with 0.1% gelatin and covered with mouse embryonic fibroblasts (MEFs), and switched to ES medium with LIF. After 3 days in culture, the blastocysts began to form outgrowths, which were subsequently disaggregated on Day 5. With extended culture, stable XEN cell lines were ultimately established around Day 30 (Figure 1B). We thus derived, using the conventional method with ES medium and LIF [21], a total of 11 XEN cell lines from 20 SCNT-derived blastocysts, at an efficiency of 55% (Table 1). For the fertilization-derived group, a very similar process of establishing XEN cell lines was observed under the same experimental conditions, and 10 XEN cell lines were finally derived from 20 fertilization-derived blastocysts, at an efficiency of 50% (Table 1). Thus, it could be concluded that NT-XENs showed an efficient derivation from blastocysts as high as that of FD-XENs.

**Table 1.** The establishment efficiency of XEN cell lines from fertilization-derived (FD) and SCNT-derived (NT) blastocysts in this study.

| Deriving type | Culture medium | Genetic background | No. of blastocysts | No. of outgrowths | No. of XEN cell lines (%) |
| --- | --- | --- | --- | --- | --- |
| FD | ES medium with LIF | B6D2F1 | 20 | 20 | 10 (50) |
| NT | ES medium with LIF | B6D2F1 | 20 | 13 | 11 (55) |
| NT | ES medium with LIF | C57BL/6J | 40 | 26 | 5 (12.5) |

Using immunofluorescence staining, we detected the expression of stem cell-specific markers in the derived NT-XENs. Like FD-XENs, the NT-XENs were able to significantly express the specific markers Gata4, Dab2, Gata6, and Sox17, but exhibited negative for the ESC-specific markers Oct4 and Nanog, and negative for the TSC-specific marker Cdx2 (Figure 2A, B, C), thereby confirming their identity of XEN cell lines.

**Figure 2.**
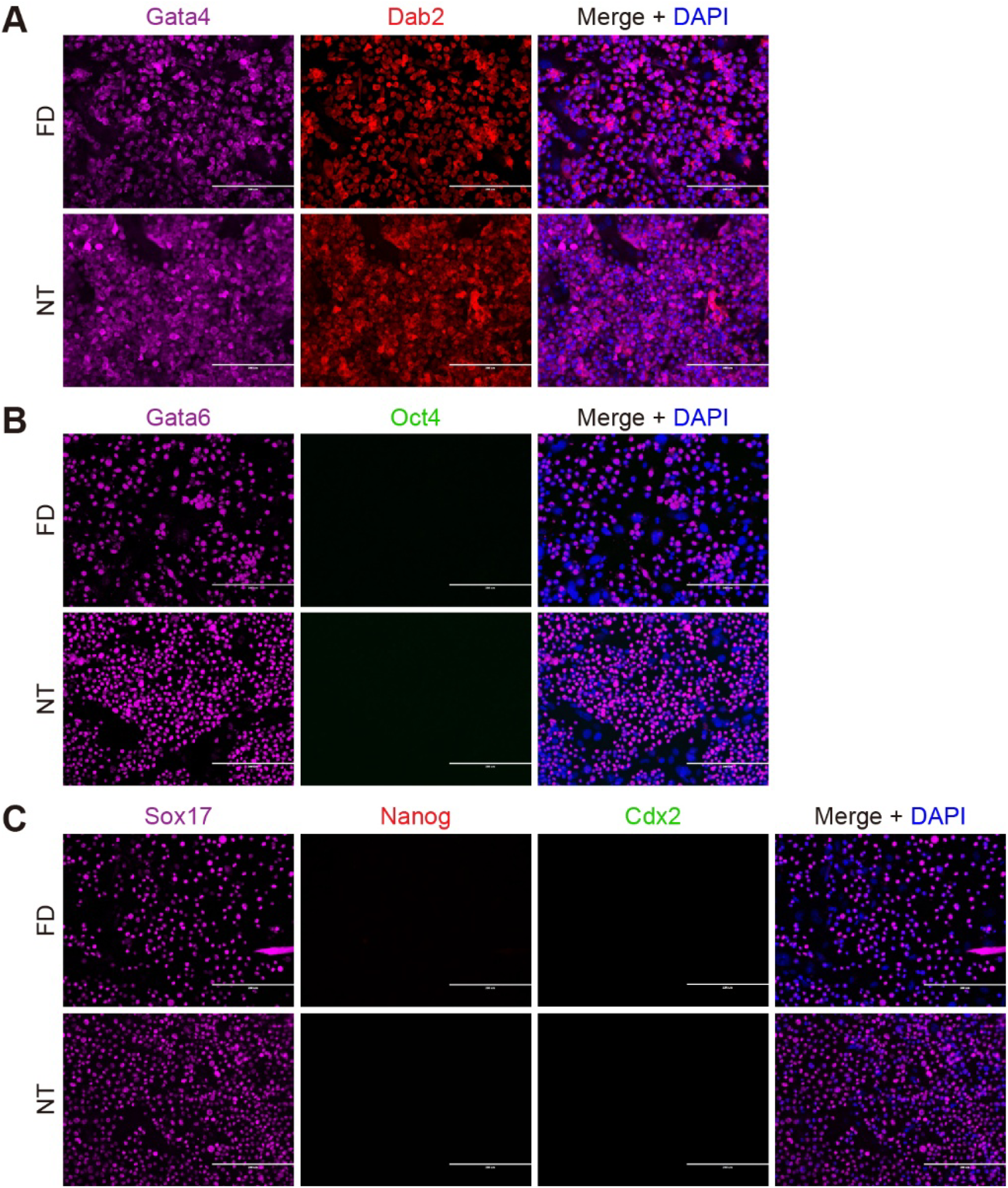
Immunofluorescence staining validates the marker protein expressions in mouse NT-XENs. The XEN-specific markers including Gata4 (A), Dab2 (A), Gata6 (B), and Sox17 (C) were strongly detected in mouse NT-XENs, whereas the ESC-specific markers Oct4 (B) and Nanog (C), and the TSC-specific marker Cdx2 (C) showed no detectable signal. Scale bar: 200 μm.

### NanoString assays validated the highly similar gene expression patterns in NT- and FD-XENs

Furthermore, we applied the NanoString multiplex platform for detecting more information on the gene expression profiles of FD-XENs and NT-XENs, respectively. A total of 222 genes, including the numerous genes with enriched expression in the early embryonic cell lineages-derived stem cell lines (e.g. ESCs, XENs, and TSCs), were detected in 4 FD-XEN lines and 6 NT-XEN lines (Additional file 1: Table S1). Inter-sample correlation clustering analysis showed that the FD-XEN and NT-XEN line samples were highly intermingled in clustering (rather than separated by group), with significant positive correlations between the replicates of the two groups (*P* < 0.001, Figure 3A), indicating that the NT-XEN lines should share a highly similar gene expression pattern with the FD-XEN lines. Of particular focus was that, similar to the FD-XEN lines, all of the NT-XEN lines presented a high abundance expression of the known XEN-specific genes (e.g. *Sox17, Sox7, Pdgfra, Gata6, Gata4*, and *Dab2*), *versus* low or no expression of ESC-specific genes (e.g. *Sox2, Oct4, Nanog, Esrrb*, and *Prdm14*) and TSC-specific genes (e.g. *Gata2, Cdx2, Elf5*, and *Tfap2c*) (Figure 3B), which thus confirmed and extended the immunofluorescence profiles. All these genetic expression characteristics further verified our high-quality derivation of XENs from SCNT-derived blastocysts.

**Figure 3.**
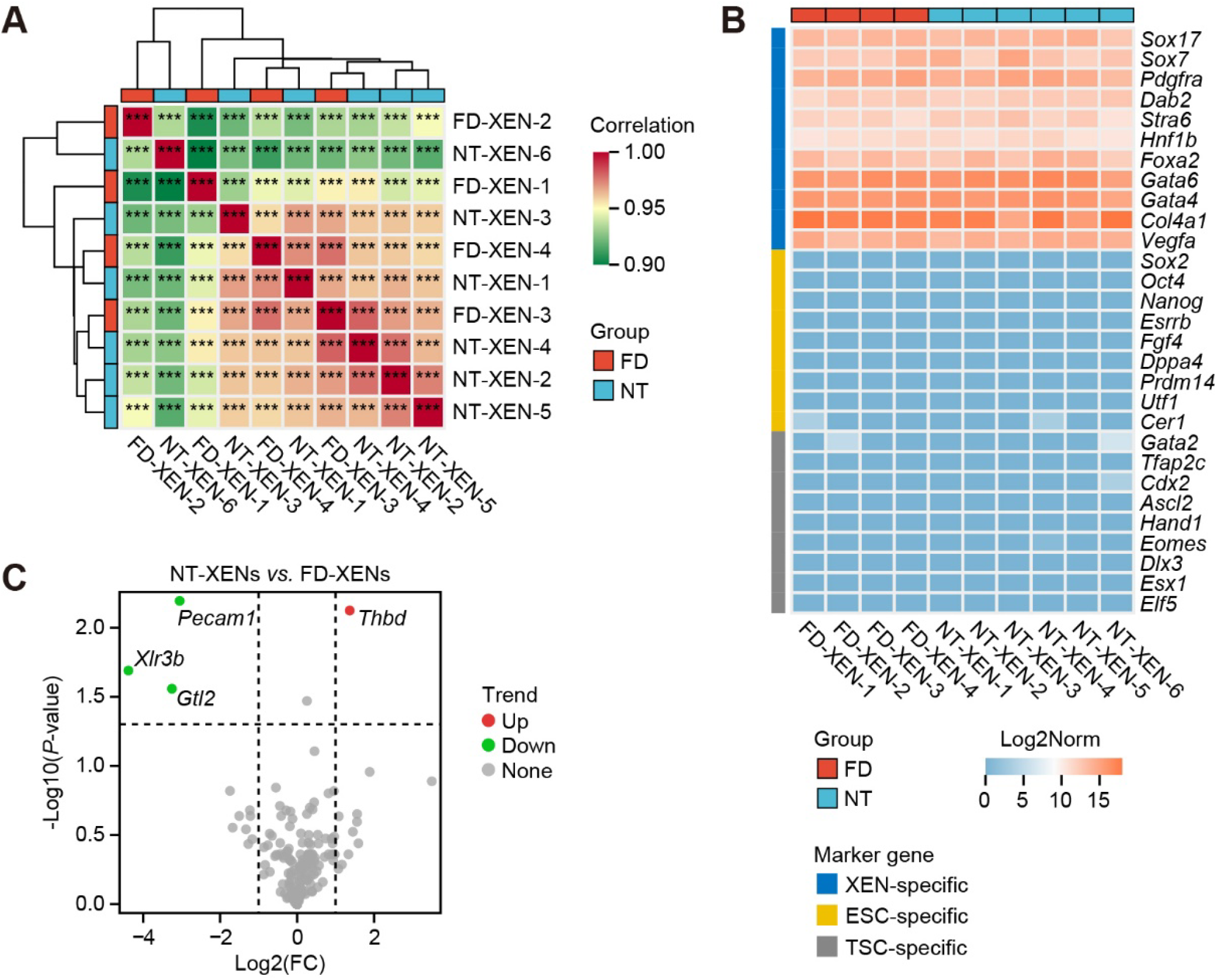
Characterization of gene expression profiles of mouse NT-XENs using NanoString. (A) Hierarchical clustering analysis of correlation among all NanoString-assayed samples (including four independent FD-XEN cell lines and six independent NT-XEN cell lines). Correlation values were calculated based on the log2-transformed normalized counts. \*\*\**P* < 0.001. (B) Expression abundance of XEN-, ESC-, and TSC-specific marker genes across all NanoString-assayed samples. The heatmap was plotted using the log2-transformed normalized counts. ESC, embryonic stem cell; TSC, trophoblast stem cell. (C) The volcano plot presents the DEG analysis result between NT-XENs and FD-XENs based on the NanoString assay data. Fold change (FC) > 2, *P*-value < 0.05.

### Minimal transcriptional divergence between FD- and NT-XENs

Based on the NanoString assay data, we performed the DEG analysis between FD-XENs and NT-XENs. As expected, the 222 surveyed genes showed virtually no differential expression overall; nevertheless, the 4 genes, *Pecam1, Thbd, Xlr3b*, and *Gtl2*, exhibited differential expression trends between the two XEN types, with *Thbd* up-regulated and the other three down-regulated in NT-XENs (fold change > 2, *P*-value < 0.05, Figure 3C, Additional file 1: Table S2). *Pecam1* is a pluripotency marker gene that is restrictedly expressed in ICM and then maintained in EPI, but not in PrE, during the second cell fate determination in blastocysts [24]. As XEN lines are the *in vitro* counterparts of PrE [21], the down-regulated *Pecam1* expression here might indicate a more differentiated state for the NT-XENs, compared with the FD-XENs. Potentially in accordance with this, another pluripotency marker gene *Gtl2*, a key maternal-allele-expressed long non-coding RNA (lncRNA) gene located in the *Dlk1*-*Dio3* imprinted domain essential for pluripotency regulation [25] and identified to exhibit restricted expression in ICM [26], was also further down-regulated in the NT-XENs. Moreover, *Thbd*, a gene preferentially expressed in the former of PE and VE [27, 28] that both derive from PrE *in vivo*, was up-regulated in the NT-XENs compared with the FD-XENs, which might imply the predicted more differentiated state of the NT-XENs towards the PE direction. *Xlr3b* is also a maternal-allele-expressed imprinted gene, which is X-linked and subject to X-chromosome inactivation (XCI) [29]. The down-regulation of *Xlr3b* expression in the NT-XENs could be possibly explained by the aberrant XCI effect that nonrandomly appears in the SCNT embryos [30] and is speculated to exist in their derivations including the NT-XENs. To summarize, the NanoString assay data revealed that the NT-XENs shared a highly similar gene expression profile with the FD-XENs, but still possessed minimal considerable transcriptional divergence from the latter.

### Lowered derivation efficiency of NT-XEN lines with inbred genetic background

Besides, we also utilized the MII oocytes and cumulus cells from C57BL/6J females to obtain reconstructed embryos, which were then cultured and seeded for deriving XEN lines under the identical experimental conditions. Eventually, we obtained only 5 XEN cell lines from 40 C57BL/6J SCNT-derived blastocysts, at a low efficiency of 12.5% (Table 1). This indicated a serious impact of the inbred genetic background on the derivation efficiency of XEN lines from SCNT-derived blastocysts, when compared with that of XEN line derivation from the SCNT-derived blastocysts with B6D2F1 hybrid genetic background.

## Discussion

In the present study, we report the first derivation of stable XEN cell lines from SCNT-derived blastocysts, providing a utilizable and renewable *in vitro* resource for investigating the reprogramming fidelity of the PrE lineage following SCNT. It is crucial, however, to acknowledge that cell lines derived *in vitro* may not perfectly represent the state of their *in vivo* counterparts [31]. The derivation process itself imposes a strong selective pressure, meaning our NT-XENs represent the outcome of both nuclear reprogramming and subsequent *in vitro* adaptation. Nevertheless, NT-XENs should be invaluable for its ability to capture and highlight the stable molecular errors in reprogramming, as studies on NT-ESCs have consistently shown that certain epigenetic aberrations, such as those affecting imprinted genes or XCI, are tenaciously retained through extensive passaging [32].

A notable finding was the high derivation efficiency of NT-XENs from hybrid B6D2F1 blastocysts (55%), which was comparable to that of fertilization-derived controls (50%). This is consistent with the reports on both NT-ESCs [19] and NT-TSCs [20], where derivation efficiencies can also rival those of conventional counterparts under optimized conditions. It should be noted that this high efficiency does not necessarily imply flawless reprogramming within the PrE of SCNT-derived blastocysts. A more plausible interpretation is that SCNT accomplishes a sufficient degree of initial reprogramming to activate the core regulatory network for PrE formation and the subsequent stringent *in vitro* culture conditions select for the most successfully reprogrammed cells to proliferate and form stable lines. In accordance with the latter, the derived NT-XENs showed highly similar transcriptional profiles with the FD-XENs, as indicated by the NanoString assay data. Thus, the high efficiency here likely reflects a combination of adequate initial reprogramming and stressful *in vitro* selection.

Despite the overall transcriptional similarity between NT-XENs and FD-XENs, the few DEGs we identified may represent key molecular signatures of incomplete reprogramming. The concurrent down-regulation of pluripotency-associated genes (*Pecam1, Gtl2*) [24, 26] and up-regulation of a PE-preferential gene (*Thbd*) [33] suggests that NT-XENs might exist in a more differentiated or lineage-restricted state. This seems to align with reports that cloned embryos can exhibit tendencies toward premature differentiation [34]. The down-regulation of *Gtl2*, a critical component of the *Dlk1*-*Dio3* imprinted domain essential for high-quality pluripotency [25, 35], further reinforces this notion. Significantly, the down-regulation of the X-linked imprinted gene *Xlr3b* is highly consistent with the aberrant XCI and subsequent misexpression of X-linked genes frequently documented in SCNT embryos [30, 36]. This finding strongly suggests that the NT-XENs should have faithfully preserved, at least partial, hallmark epigenetic errors of the SCNT origin, validating their utility as a model for studying such defects.

## Conclusions

Our study successfully establishes and characterizes NT-XEN cell lines, demonstrating that while they are remarkably similar to their fertilization-derived counterparts, they harbor subtle but significant transcriptional differences that likely reflect the molecular legacy of their SCNT origin. The establishment of these cell lines fills a critical gap in the field, providing an important and previously unavailable resource for exploring the molecular signatures causally linked to the yolk sac abnormalities and developmental failure endemic to cloned embryos.

## Declarations

### Data availability

All data generated or analyzed during this study are included in this published article and its supplementary information files. The processed and analyzed NanoString gene expression data are available in Additional file 1.

## Acknowledgements

We sincerely thank Dr. Han Gao, a Ph.D. candidate from the Kunming Institute of Zoology, Chinese Academy of Sciences, for her help with the experimental data processing.

## Authors’ contributions

WB and JL conceived this study. SL and SW designed the experiments. SL, SW, GL and MH performed the experiments. WB and JL drew the diagrams and drafted the manuscript. All authors read and approved the final version of the manuscript.

## Supplementary materials

Additional file 1: Table S1, S2.

## References

1. Hochedlinger K, Jaenisch R: Nuclear transplantation, embryonic stem cells, and the potential for cell therapy. The New England journal of medicine 2003, 349(3):275–286.

2. Han YM, Kang YK, Koo DB, Lee KK: Nuclear reprogramming of cloned embryos produced in vitro. Theriogenology 2003, 59(1):33–44.

3. Rideout WM, 3rd, Eggan K, Jaenisch R: Nuclear cloning and epigenetic reprogramming of the genome. Science 2001, 293(5532):1093–1098.

4. Yang X, Smith SL, Tian XC, Lewin HA, Renard JP, Wakayama T: Nuclear reprogramming of cloned embryos and its implications for therapeutic cloning. Nat Genet 2007, 39(3):295–302.

5. Tanaka S, Oda M, Toyoshima Y, Wakayama T, Tanaka M, Yoshida N, Hattori N, Ohgane J, Yanagimachi R, Shiota K: Placentomegaly in cloned mouse concepti caused by expansion of the spongiotrophoblast layer. Biology of reproduction 2001, 65(6):1813–1821.

6. Ogura A, Inoue K, Wakayama T: Recent advancements in cloning by somatic cell nuclear transfer. Philosophical transactions of the Royal Society of London Series B, Biological sciences 2013, 368(1609):20110329.

7. Nishioka N, Inoue K, Adachi K, Kiyonari H, Ota M, Ralston A, Yabuta N, Hirahara S, Stephenson RO, Ogonuki N et al: The Hippo signaling pathway components Lats and Yap pattern Tead4 activity to distinguish mouse trophectoderm from inner cell mass. Dev Cell 2009, 16(3):398–410.

8. Strumpf D, Mao CA, Yamanaka Y, Ralston A, Chawengsaksophak K, Beck F, Rossant J: Cdx2 is required for correct cell fate specification and differentiation of trophectoderm in the mouse blastocyst. Development 2005, 132(9):2093–2102.

9. Yamanaka Y, Lanner F, Rossant J: FGF signal-dependent segregation of primitive endoderm and epiblast in the mouse blastocyst. Development 2010, 137(5):715–724.

10. Smyth N, Vatansever HS, Murray P, Meyer M, Frie C, Paulsson M, Edgar D: Absence of basement membranes after targeting the LAMC1 gene results in embryonic lethality due to failure of endoderm differentiation. The Journal of cell biology 1999, 144(1):151–160.

11. Brennan J, Lu CC, Norris DP, Rodriguez TA, Beddington RS, Robertson EJ: Nodal signalling in the epiblast patterns the early mouse embryo. Nature 2001, 411(6840):965–969.

12. Rossant J, Cross JC: Placental development: lessons from mouse mutants. Nat Rev Genet 2001, 2(7):538–548.

13. Beddington RS, Robertson EJ: Axis development and early asymmetry in mammals. Cell 1999, 96(2):195–209.

14. Evans MJ, Kaufman MH: Establishment in culture of pluripotential cells from mouse embryos. Nature 1981, 292(5819):154–156.

15. Kunath T, Arnaud D, Uy GD, Okamoto I, Chureau C, Yamanaka Y, Heard E, Gardner RL, Avner P, Rossant J: Imprinted X-inactivation in extra-embryonic endoderm cell lines from mouse blastocysts. Development 2005, 132(7):1649–1661.

16. Tanaka S, Kunath T, Hadjantonakis AK, Nagy A, Rossant J: Promotion of trophoblast stem cell proliferation by FGF4. Science 1998, 282(5396):2072–2075.

17. Artus J, Piliszek A, Hadjantonakis AK: The primitive endoderm lineage of the mouse blastocyst: sequential transcription factor activation and regulation of differentiation by Sox17. Dev Biol 2011, 350(2):393–404.

18. Wakayama T, Tabar V, Rodriguez I, Perry AC, Studer L, Mombaerts P: Differentiation of embryonic stem cell lines generated from adult somatic cells by nuclear transfer. Science 2001, 292(5517):740–743.

19. Tachibana M, Amato P, Sparman M, Gutierrez NM, Tippner-Hedges R, Ma H, Kang E, Fulati A, Lee HS, Sritanaudomchai H et al: Human embryonic stem cells derived by somatic cell nuclear transfer. Cell 2013, 153(6):1228–1238.

20. Oda M, Tanaka S, Yamazaki Y, Ohta H, Iwatani M, Suzuki M, Ohgane J, Hattori N, Yanagimachi R, Wakayama T et al: Establishment of trophoblast stem cell lines from somatic cell nuclear-transferred embryos. Proc Natl Acad Sci U S A 2009, 106(38):16293–16297.

21. Niakan KK, Schrode N, Cho LT, Hadjantonakis AK: Derivation of extraembryonic endoderm stem (XEN) cells from mouse embryos and embryonic stem cells. Nature protocols 2013, 8(6):1028–1041.

22. Lin J: Derivation of mouse extraembryonic endoderm stem cell lines, and exclusive transmission of the embryonic stem cell-derived genome through the mouse germline. Doctoral dissertation. Frankfurt am Main: Johann Wolfgang Goethe-Universität in Frankfurt am Main; 2018.

23. Jiang S, Li H, Zhang L, Mu W, Zhang Y, Chen T, Wu J, Tang H, Zheng S, Liu Y et al: Generic Diagramming Platform (GDP): a comprehensive database of high-quality biomedical graphics. Nucleic Acids Res 2025, 53(D1):D1670–d1676.

24. Robson P, Stein P, Zhou B, Schultz RM, Baldwin HS: Inner cell mass-specific expression of a cell adhesion molecule (PECAM-1/CD31) in the mouse blastocyst. Dev Biol 2001, 234(2):317–329.

25. Stadtfeld M, Apostolou E, Akutsu H, Fukuda A, Follett P, Natesan S, Kono T, Shioda T, Hochedlinger K: Aberrant silencing of imprinted genes on chromosome 12qF1 in mouse induced pluripotent stem cells. Nature 2010, 465(7295):175–181.

26. Han Z, Yu C, Tian Y, Zeng T, Cui W, Mager J, Wu Q: Expression patterns of long noncoding RNAs from Dlk1-Dio3 imprinted region and the potential mechanisms of Gtl2 activation during blastocyst development. Biochemical and biophysical research communications 2015, 463(3):167–173.

27. Smith MK, Clark CC, McCoski SR: Technical note: improving the efficiency of generating bovine extraembryonic endoderm cells. Journal of animal science 2020, 98(7).

28. Yagi S, Tagawa Y, Shiojiri N: Transdifferentiation of mouse visceral yolk sac cells into parietal yolk sac cells in vitro. Biochemical and biophysical research communications 2016, 470(4):917–923.

29. Raefski AS, O’Neill MJ: Identification of a cluster of X-linked imprinted genes in mice. Nat Genet 2005, 37(6):620–624.

30. Inoue K, Kohda T, Sugimoto M, Sado T, Ogonuki N, Matoba S, Shiura H, Ikeda R, Mochida K, Fujii T et al: Impeding Xist expression from the active X chromosome improves mouse somatic cell nuclear transfer. Science 2010, 330(6003):496–499.

31. Nichols J, Smith A: Naive and primed pluripotent states. Cell Stem Cell 2009, 4(6):487–492.

32. Humpherys D, Eggan K, Akutsu H, Hochedlinger K, Rideout WM, 3rd, Biniszkiewicz D, Yanagimachi R, Jaenisch R: Epigenetic instability in ES cells and cloned mice. Science 2001, 293(5527):95–97.

33. Weiler-Guettler H, Aird WC, Rayburn H, Husain M, Rosenberg RD: Developmentally regulated gene expression of thrombomodulin in postimplantation mouse embryos. Development 1996, 122(7):2271–2281.

34. Boiani M, Eckardt S, Schöler HR, McLaughlin KJ: Oct4 distribution and level in mouse clones: consequences for pluripotency. Genes Dev 2002, 16(10):1209–1219.

35. Liu L, Luo GZ, Yang W, Zhao X, Zheng Q, Lv Z, Li W, Wu HJ, Wang L, Wang XJ et al: Activation of the imprinted Dlk1-Dio3 region correlates with pluripotency levels of mouse stem cells. J Biol Chem 2010, 285(25):19483–19490.

36. Eggan K, Akutsu H, Hochedlinger K, Rideout W, 3rd, Yanagimachi R, Jaenisch R: X-Chromosome inactivation in cloned mouse embryos. Science 2000, 290(5496):1578–1581.

